# Antiviral and Neuroprotective Abilities of Influenza Virus Infection in Tractable Brain Organoids

**DOI:** 10.1101/2022.03.02.482634

**Authors:** Xiaodong Zhang, Haishuang Lin, Liangzhen Dong, Qing Xia

**Author notes:** These authors contributed equally to this work (Xiaodong Zhang, Haishuang Lin, Liangzhen Dong).

## Abstract

Human pluripotent stem cell (hPSC)-derived brain organoids offer an unprecedented opportunity for various applications as an *in vitro* model, such as modeling virus infection and drug screening. In this study, we present an experimental brain organoid platform for modeling infection with multiple viruses (e.g., influenza virus or enterovirus). Brain organoids challenged by influenza viruses (H1N1-WSN and H3N2-HKT68) had decreased overall organoid size, similar to ZIKA virus infection, while enteroviruses (EV68 and EV71) infected brain organoids displayed the opposite result. Then, we studied the molecular events in WSN-infected organoids, and we found that WSN could widely infect multiple cell types, and preferentially infected MAP2+ neurons compared to SOX2+ neural stem cells (NSCs) and GFAP+ astrocytes in brain organoids, and induced apoptosis of NSCs and neurons, but not astrocytes. The inflammatory responses in organoids observed to occur (Tumor necrosis factor alpha, interferon gamma, and interleukin 6) after WSN infection may further facilitate brain damage. Furthermore, transcriptional profiling revealed several upregulated genes (*CSAG3* and *OAS2*) and downregulated genes (*CDC20B, KCNJ13, OTX2-AS1, CROCC2*, and *F5*) after WSN infection for 24 hpi and 96 hpi, implicating antiviral drugs development responses to WSN. Finally, we explored neurotrophic factors (e.g., BDNF, GDNF, and NT3) and PYC-12 as antiviral and neuroprotective reagents, which could significantly suppress virus infection, apoptosis, and inflammatory responses. Collectively, we established a tractable experimental model system to investigate the impact and mechanism of virus infection on human brain development, and provide a platform for rapidly screening therapeutic compounds, advancing the development of antiviral strategies.

## Introduction

Central nervous system (CNS) infections are one of the most critical public health problems, and they lead to high morbidity and mortality every year, predominately in young children, the elderly, and the immunocompromised (Englund et al., 2011; Keipp Talbot and Falsey, 2010; Tregoning and Schwarze, 2010). CNS infections often lead to neurological sequelae, including epilepsy (Vehapoglu et al., 2015), and neurodevelopmental disorders, such as attention-deficit/hyperactivity disorder (ADHD) (Hoekstra, 2019) and autism spectrum disorder (ASD) (Sauer et al., 2021). Some known neurotropic viruses, such as measles virus (MV), herpes virus, and human immunodeficiency virus (HIV), can cause CNS infections (Koyuncu et al., 2013). Moreover, respiratory viruses including human respiratory syncytial virus (HRSV), influenza virus (IV), and coronavirus (Cov) have also become key factors responsible for CNS pathologies. In particular, COVID-19, which is caused by severe acute respiratory syndrome coronavirus 2 (SARS-CoV-2), has been characterized by respiratory failure in critically ill patients. The outbreak of COVID-19 has led to a pandemic, and serious and even fatal manifestations have been seen in the brain (Johansson et al., 2020). The annual pandemic of influenza A virus (IAV) has been shown to cause neurodegenerative diseases in clinical investigations (McGavern and Kang, 2011). IAV is often the causative agent of upper respiratory infection by recognizing sialic acids (SA) receptor in humans affecting people of all ages (Han et al., 2018). Although the influenza virus primarily infects the lungs, its neuropathological effects have also been shown in the clinic, including febrile seizures, myelitis, focal encephalitis, and even meningitis (Liang et al., 2018). However, the neuropathogenesis of these manifestations remains elusive, owing to the lack of physiological *in vitro* models instead of patient-derived brain tissues. Therefore, there is an urgent need of *in vitro* physiological models of virus infection to identify the neuropathological mechanism in the brain.

Human pluripotent stem cell (hPSC)-derived organoids, which are an *in vitro* self-organizing and self-renewing three-dimensional tissue recapitulating the cytoarchitecture and functional components of human tissue, can be amendable to tissue development, disease modeling, drug screening, and therapeutic discovery (De Crignis et al., 2021; Qian et al., 2019). For example, Chen et al. (2021) generated serum-exposed brain organoids that were able to recapitulate Alzheimer’s disease (AD)-like pathologies, providing a powerful platform for both mechanistic study and therapeutic development of AD treatments. Dang et al. (2016) utilized human cerebral organoids to demonstrate that Zika virus (ZIKV) could decrease the number of neural progenitors through activation of the innate immune receptor TLR3, resulting in dysregulation of a network of genes involved in axon guidance, neurogenesis, differentiation, and apoptosis. In addition, Clevers’ group leveraged colorectal cancer (CRC) organoid lines to screen target-known inhibitors and chemotherapy drugs (Van De Wetering et al., 2015), and also used breast cancer organoid lines to screen EGFR/AKT/mTORC pathway inhibitors (Sachs et al., 2018). Currently, several organoids (e.g., lung, liver, gut, brain, and kidney) mimicking different tissues have been used to study the effects of SARS-CoV-2 on tissues *in vitro*, and tissue histopathology (Yu, 2021). The advantage of brain organoids over cell lines is that organoids model the three-dimensional organ to achieve multiple cellular interactions, and rather than animal models, the use of organoids enables studies of humans. Therefore, organoid technology offers a significant advantage in terms of mimicking *in vivo* tissue structures for testing antiviral compounds. To date, influenza virus-infected brain organoid use to identify neuropathogenesis or for drug screening has not been reported. In this study, we evaluated the potential of brain organoids as *in vitro* infection models, and studied the neuropathogenesis of brain organoids challenged by WSN using hPSC-derived brain organoids. We found that WSN preferentially infected MAP2+ neurons in brain organoids, and further damaged the brain by eliciting inflammatory factor release (TNF-α, INF-γ, and IL-6), as well as inducing the apoptosis of NSCs and neurons, but not astrocytes. Additionally, RNA transcriptomic profiles revealed new protein targets and noncoding RNAs for the development of antiviral strategies. Finally, we conducted an antiviral study and found that some specific neurotrophic factors (e.g., BDNF, GDNF, and NT3) and a potential antiviral drug candidate, PYC-12, could both significantly decrease virus replication to achieve antiviral and neuroprotective effects. Collectively, human organoids were shown to serve as an invaluable tool in the field of virus research to investigate the molecular mechanisms underlying virus replication and antiviral drug screening.

## Results

### Generation and characterization of cerebral organoids from hPSCs

To study the neuropathogenesis of virus-infected brain organoids instead of patient-derived brain tissue, we first generated human pluripotent stem cell (hPSC)-derived brain organoids as previously described (Lancaster and Knoblich, 2014), with slight modifications. Briefly, hPSCs were first dissociated into single cells, followed by centrifugation at 100 g for 3 minutes. hPSCs were allowed to form embryoid bodies (EBs) in Essential 8 medium containing 10 µM ROCK inhibitor for 4 days, incubated in neural induction medium to develop neuroectoderm, and then transferred to a Matrigel droplet. Finally, these Matrigel droplet encapsulated tissues were transferred to a spinning bioreactor, which promoted nutrient and oxygen exchange to further develop into brain organoids (**Fig. 1A**). The representative bright field images display the entire development of brain organoids at indicated time points (**Fig. 1B**). In the process of organoid development, the diameter (µm) of organoids increased, while the area (µm^2^) increased in early stages and decreased obviously in later stages (**Fig. 1C, D**). Next, we performed immunostaining of brain organoids at days 15, 30, 60, and 120 with the neural stem cell marker-SOX2 and the neuron marker-MAP2 (**Fig. 1E, F**). Statistical analysis showed that the majority of the cells on day 15 and day 30 organoids were neural stem cells (SOX2+), while the majority of the cells on day 60 and day 120 organoids were neurons (MAP2+) (**Fig. 1G**). Finally, quantitative real-time polymerase chain reaction (RT-PCR) confirmed the specific markers in the entire process of brain organoid development, including the hPSC markers, *OCT4* and *NANOG*; the NSC marker, *PAX6*; the forebrain markers, *FOXG1* and *SIX3*; the hindbrain markers, *KROX20* and *ISL1*; the neuron marker, *TUJ1*; and the deep-layer cortical neuron markers, *CTIP2* and *TBR1* (**Fig. 1H**). In summary, these results suggested that there was robust differentiation of brain organoids and following applications of virus infection model.

**Figure 1.**
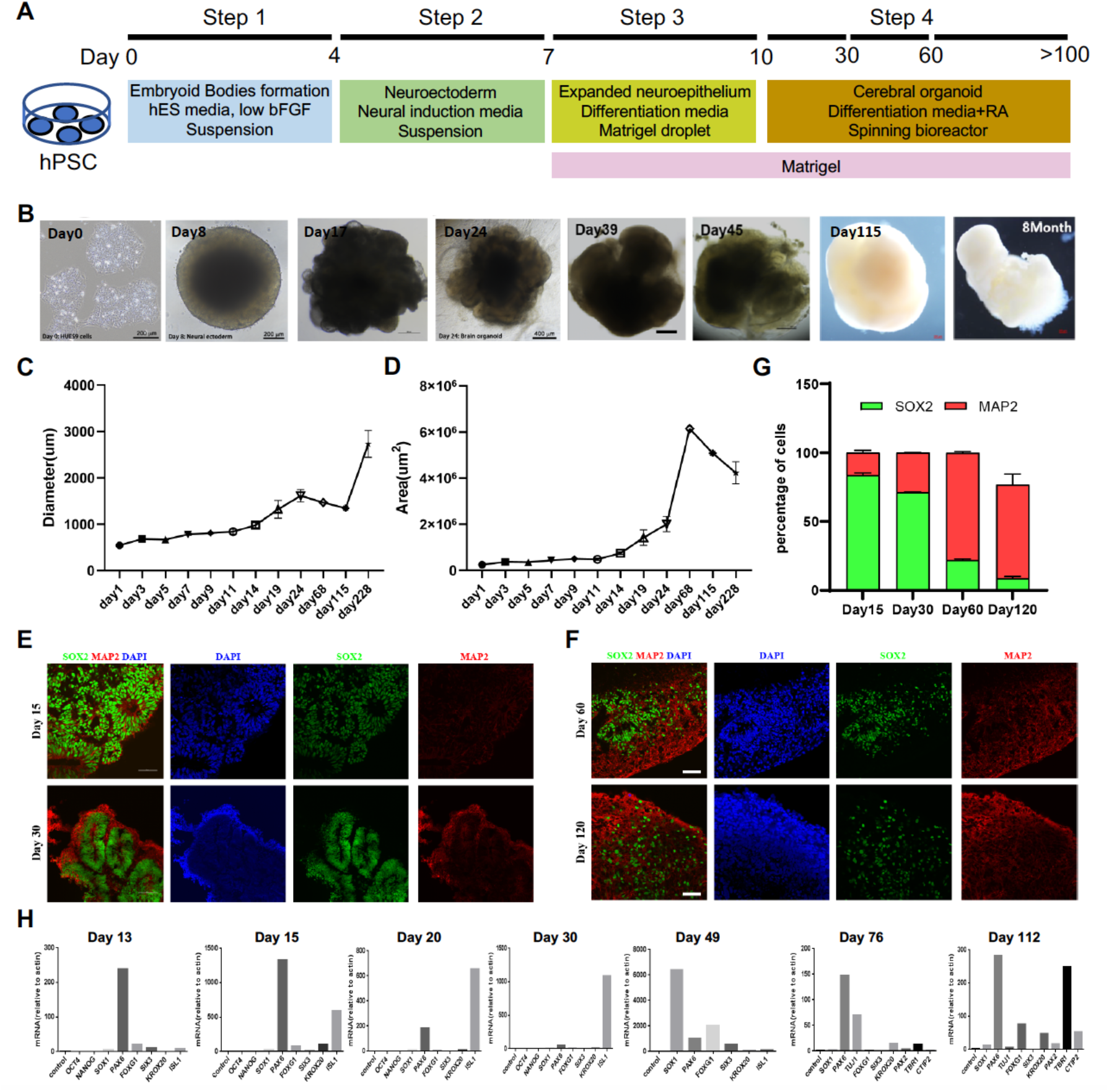
Generation and characterization of brain organoids. (**A**) Schematic illustration of organoid generation. (**B**) Representative bright-field images of brain organoids derived from hPSCs at day 0, day 8, day 17, day 24, day 39, day 45, day 115, and 8 months. Scale bars, 200 µm and 400 µm. (**C, D**) Growth kinetics of area (µm^2^) and diameter (µm) of brain organoids during development. (**E, F**) Immunostaining of brain organoids with SOX2+ NSCs and MAP2+ neurons on day 15, 30, 60 and 120. Scale bars, 100 µm. (**G**) The percentage of SOX2+ and MAP2+ cells in day 15, 30, 60, and 120 brain organoids. The majority of cells on days 15 and 30 organoids were SOX2+ neural stem cells and MAP2+ neurons on days 60 and 120 organoids. (**H**) Quantitative RT-PCR analysis of the specific markers of brain organoids at indicated timepoints. hPSC markers: *OCT4* and *NANOG*; NSC marker: *PAX6*; forebrain markers: *FOXG1* and *SIX3*; hindbrain markers: *KROX20* and *ISL1*; neuron marker: *TUJ1*; deep-layer cortical neuron markers: *CTIP2* and *TBR1*.

### Brain organoid as an in vitro model for distinct virus infection

To model the effects of virus infection on early human brain development, we used hPSC-derived brain organoids at day 40 of differentiation. We challenged these day 40 brain organoids with a diverse panel of viruses, including influenza viruses (H1N1-WSN and H3N2-HKT68), enteroviruses (EV68 and EV71) and severe fever with thrombocytopenia syndrome virus (SFTSV) at indicated timepoints, and identified the optimal doses for virus infection. These were 1×10^6^ pfu for WSN, 5×10^6^ pfu for H3N2, 8×10^4^ pfu for EV68, and 4×10^6^ pfu for EV71 (**Fig. 2A**). The representative bright field images show a decrease in overall organoid size in the influenza virus infection group at indicated virus doses (**Fig. 2B, D**), which was similar to Zika virus infection but not as severe (Tang et al., 2016). Statistical analysis of diameters (µm) and areas (µm^2^) for influenza virus-infected organoids showed a dose-dependent size decrease (**Fig. 2C, E**). Surprisingly, when we challenged day 40 organoids with the other two H1N1 subtypes, PR8 and CA07, we observed a slight increase of organoid size at indicated virus doses through representative bright field images and statistical analysis of diameters (µm) and areas (µm^2^) (**Fig. S1**). Meanwhile, enteroviruses (EV68 and EV71)-infected brain organoids resulted in a significant increase of overall organoid size at indicated virus doses through bright field images and statistical analysis of their diameters and areas (**Fig. 2F–I**). However, SFTSV-infected brain organoids showed no obvious changes in overall organoid size (**Fig. 2J, K**). Therefore, the morphological changes of organoids challenged by multiple viruses may be different due to different mechanisms.

**Figure 2.**
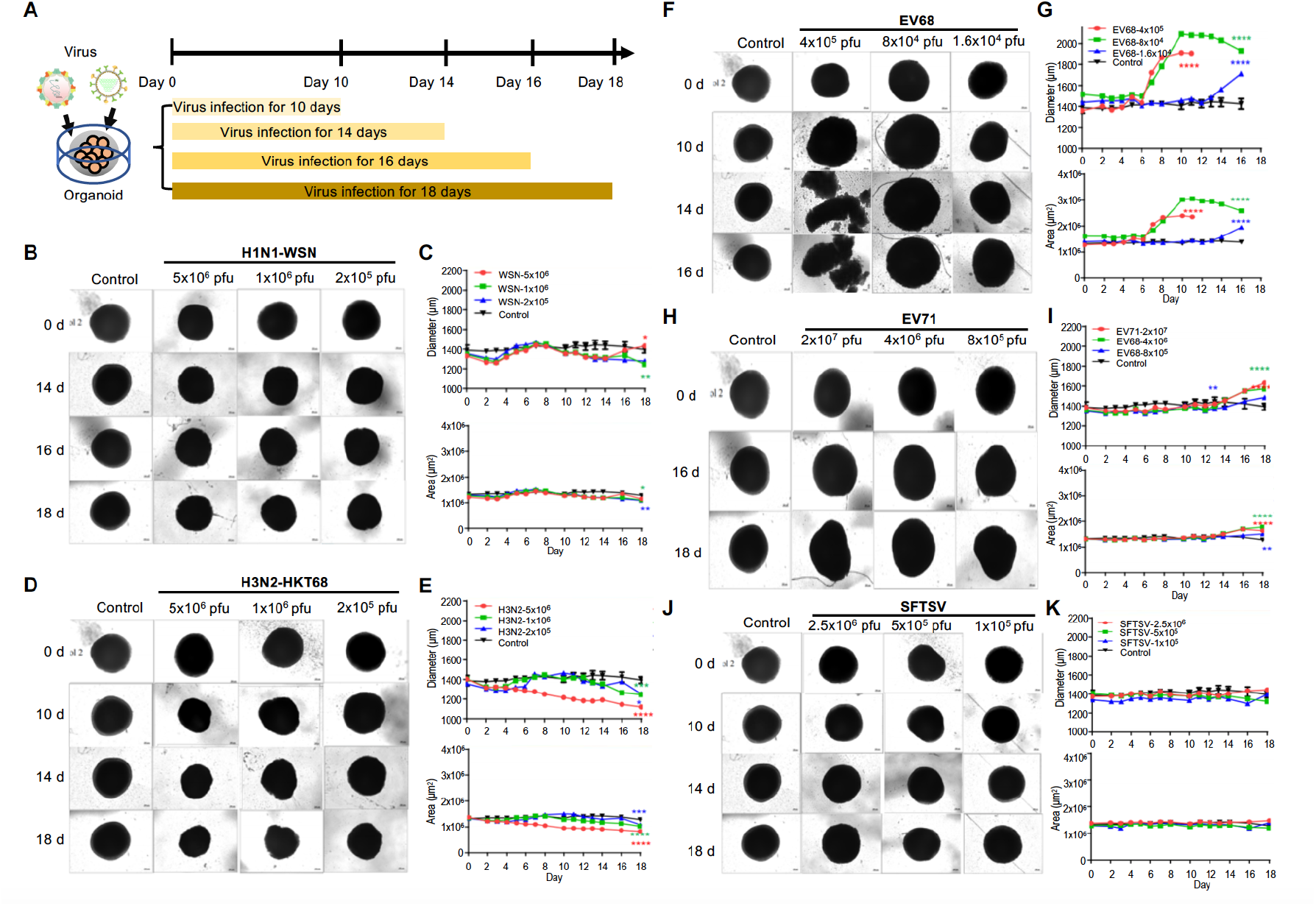
Organoids as virus infection models. (**A**) Schematic illustration of experimental design. Day 40 brain organoids were infected with viruses at indicated times. (**B, D, F, H, J**) Representative bright-field images of day 40-brain organoids infected with viruses at indicated time points, including H1N1-WSN, H3N2-HKT68, EV68, EV71, and SFTSV. The infected concentrations for H1N1-WSN were 5×10^6^ pfu, 1×10^6^ pfu, and 2×10^5^ pfu; for H3N2, they were 5×10^6^ pfu, 1×10^6^ pfu, and 2×10^6^ pfu; for EV68, they were 4×10^5^ pfu, 8×10^4^ pfu, and 1.6×10^4^ pfu; for EV71, they were 2×10^7^ pfu, 4×10^6^ pfu, and 8×10^5^ pfu. Scale bars, 50 µm. (**C, E, G, I, K**) Statistical analysis of area (µm^2^) and diameter (µm) of brain organoid infected with viruses at indicated time points.

Next, to better understand the infection mechanism of brain organoids with virus, we primarily focused on the study of WSN infected brain organoids. These brain organoids were generated as mentioned previously and infected on days 30, 60, and 250 (**Fig. 3A**). Before WSN infection of organoids, we first confirmed that ~20% NESTIN+ NSCs derived from hPSCs and ~10% GFAP+ astrocyte could be infected by WSN at the cellular level (**Fig. 3B**). The advantage of brain organoids over cell lines is that organoids model a three-dimensional organ to achieve multiple cellular interactions, and the advantage of organoids over animal models is that organoids enable studies of humans. Immunostaining of day 30 and day 60 brain organoids with SOX2+ NSCs and MAP2+ neurons showed a time-dependent increase of NP+ cells compared to mock infection, especially for MAP2+ neurons (**Fig. 3C, D**). The statistical percentage of NP+ cells demonstrated that WSN was more likely to infect MAP2+ neurons (~60%) at 4 dpi, while only ~10% – 20% of NSCs were infected at 1 dpi and 4 dpi, respectively (**Fig. 3E, H**). The intracellular and extracellular virus titers showed a time-dependent increase after WNS infection in day 30 or day 60 organoids (**Fig. 3F, G, I, J**). To further investigate long-term infection, we cultured brain organoids for 250 days. Immunostaining of day 250 organoids revealed that ~30% of SOX2+ NSCs, ~60% of MAP2+ neurons, and ~10% of GFAP+ astrocytes were infected by WSN (**Fig. 3K, L**). The extracellular virus titers showed a time-dependent increase in WSN infected day 250 organoids (**Fig. 3M**). These results suggested that MAP2+ neurons were more susceptible to WSN infection compared to NSCs or astrocytes in brain organoids (**Fig. 3N**).

**Figure 3.**
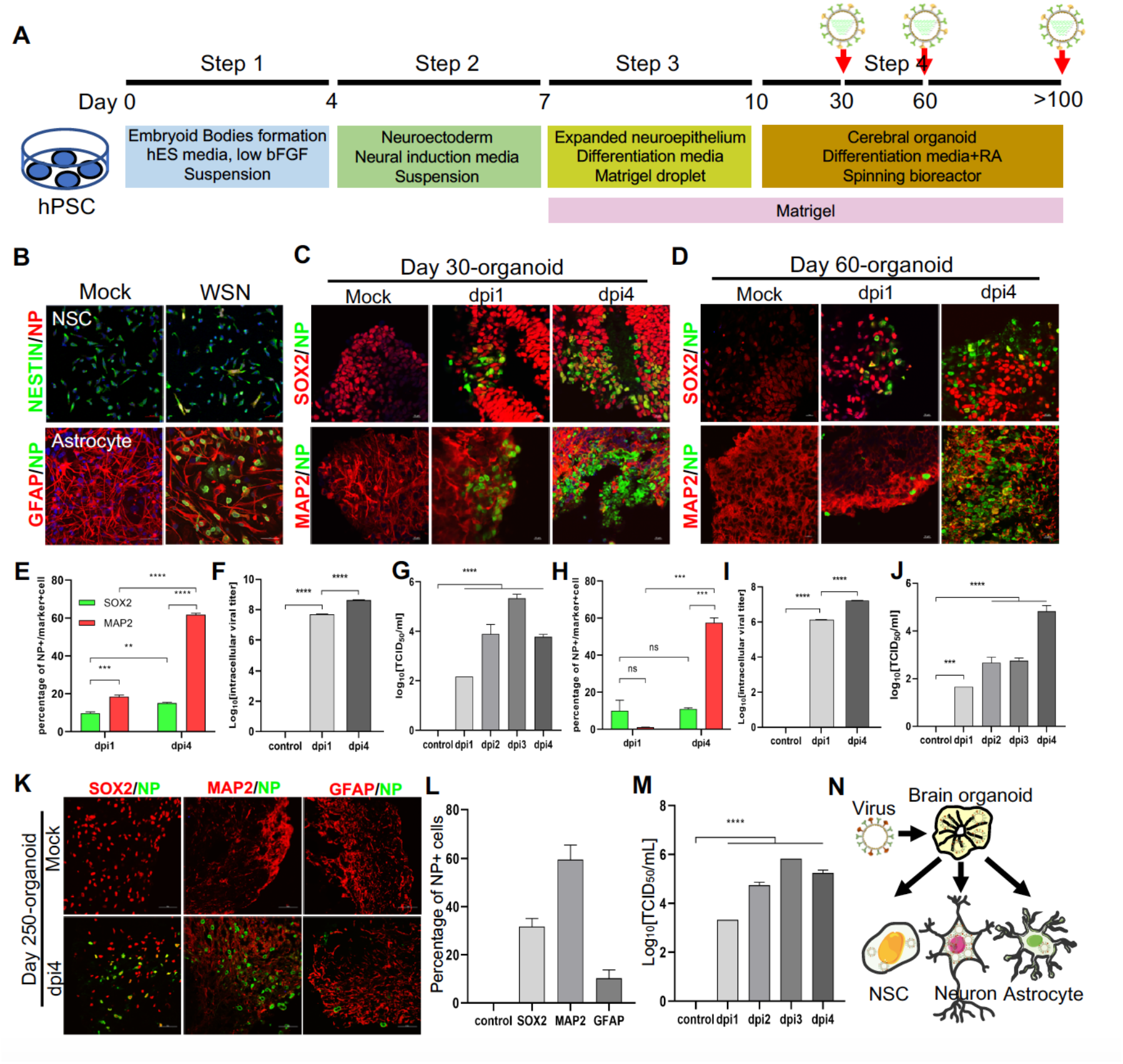
Modeling influenza virus infection *in vitro* using brain organoids. (**A**) Schematic illustration of the experimental design of brain organoids infected with WSN at indicated time points. (**B**) Immunostaining of neural stem cells and astrocytes infected with WSN. Scale bars, 50 µm. (**C, D**) Immunostaining of neural stem cells and neurons in day 30 and day 60 brain organoids infected with WSN for 1 day and 4 days. Scale bars, 100 µm. (**E, H**) The percentage of NP+ cells in infected day 30 and day 60 brain organoids. ~10% of SOX2+ NSCs and ~60% of MAP2+ neurons were infected with WSN. (**F, G, I, J**) The viral titers of intracellular and supernatants in day 30 and day 60 brain organoids. (**K**) Immunostaining of neural stem cells, neurons and astrocytes of day 250 brain organoids infected with WSN for 1 day and 4 days. Scale bars, 100 µm. (**L**) The percentage of NP+ cells in infected brain organoids. ~30% of SOX2+ NSCs, ~60% of MAP2+ neurons and 10% astrocytes were infected with WSN. (**M**) The viral titers of supernatants on day 250 brain organoids. (**N**) Schematic illustration of WSN preferentially infected neurons in brain organoids.

### Transcriptomic profiling of WSN infected organoids

To investigate the underlying molecular mechanisms at the gene level, we performed transcriptomic profiling of WSN-infected organoids at 1 dpi and 4 dpi compared to mock-infected organoids. Hierarchical clustering analysis showed the significant differences in gene expression between 1 dpi and 4 dpi organoids compared to mock-infected organoids (**Fig. 4A**). A Venn diagram shows that there were more gene alterations at 4 dpi compared to 1 dpi, and 266 genes were upregulated while 117 genes were downregulated in both groups (**Fig. 4B**). GO terms were mainly enriched in the regulation of cellular metabolic process, negative regulation of developmental process, and innate immune response at 1 dpi, while they were mainly enriched in negative regulation of nervous system development, and neuron differentiation at 4 dpi (**Fig. 4C**). KEGG signaling pathway terms were enriched in virus infection pathways at 1 dpi, while they were enriched in MAPK signaling pathway and glycolysis/gluconeogenesis at 4 dpi (**Fig. S2A**). Next, we performed a detailed gene expression analysis at 1 dpi and 4 dpi. In the top 10 of upregulated and downregulated genes, *FABP1, CDX2, FGG, ISX, SI, GSTA1, GBP1P1, CXCL10, KRT20*, and *CXCL11* were significantly upregulated, while *MCIDAS, MFRP, CCKAR, CDC20B, AL355812*.*1, SLC39A12, CCNO, ABCA4, KCNJ13*, and *OTX2-AS1* were significantly downregulated in the 1 dpi group. *CCL7, HIST1H3PS1, RN7SL472P, NCOA4P2, WDR95P, IL6, ZSCAN4, AP001331*.*1, AC108134*.*2*, and *GBP1P1* were significantly upregulated, while *DAPL1, MIR217HG, SIX6, AL451127*.*1, TFAP2D, AC016044*.*1, SIX3OS1_2, VSX1, CLRN1*, and *AL138826*.*1* were significantly downregulated in the 4 dpi group (**Fig. 4D**). *GBP1P1, CXCL10, CXCL11, CCL7, CSAG3, OAS2*, and *NCOA4P2* were upregulated and *CDC20B, AL355812*.*1, KCNJ13, OTX2-AS1, CROCC2*, and *F5* were downregulated in both groups (**Fig. 4E**). In most cells, interferon (IFN) response is a major first line of defense against viral infection. Viral infection triggers the production of IFNs, which then bind to ubiquitously expressed receptors on nearby cells and induce a powerful transcriptional program comprised hundreds of antiviral IFN-stimulated genes (ISGs) (Malterer et al., 2014). In this study, hierarchical clustering analysis identified a set of ISGs (e.g., *IFITM1/2/3, BST2*, and *SLC16A1*) that were highly induced in the early stages of WSN infection (1 dpi), while significantly decreased in late stages of infection (4 dpi) (**Fig. 4F**). This indicated that ISGs played the crucial role against viral invasion during early infection. We also analyzed some DEGs of transcription factors (**Fig. S2B**), inflammatory factors (**Fig. S2C**), and metabolic genes (**Fig. S2D**) that were associated with virus infection. We found that they showed significant changes between 1 dpi and 4 dpi compared to mock infection. Moreover, a protein-protein interaction network of WSN infected brain organoids at 4 dpi was more robust than at 1 dpi (**Fig. S2E**). In addition, WSN-infected organoids caused not only protein gene changes, but also some noncoding RNA levels, such as *BISPR* and *MIR4435-2HG* (**Fig. S3**). Collectively, these results suggested that WSN infection of brain organoids resulted in considerable gene alterations, and implicating these genes for the development of new antiviral strategies.

**Figure 4.**
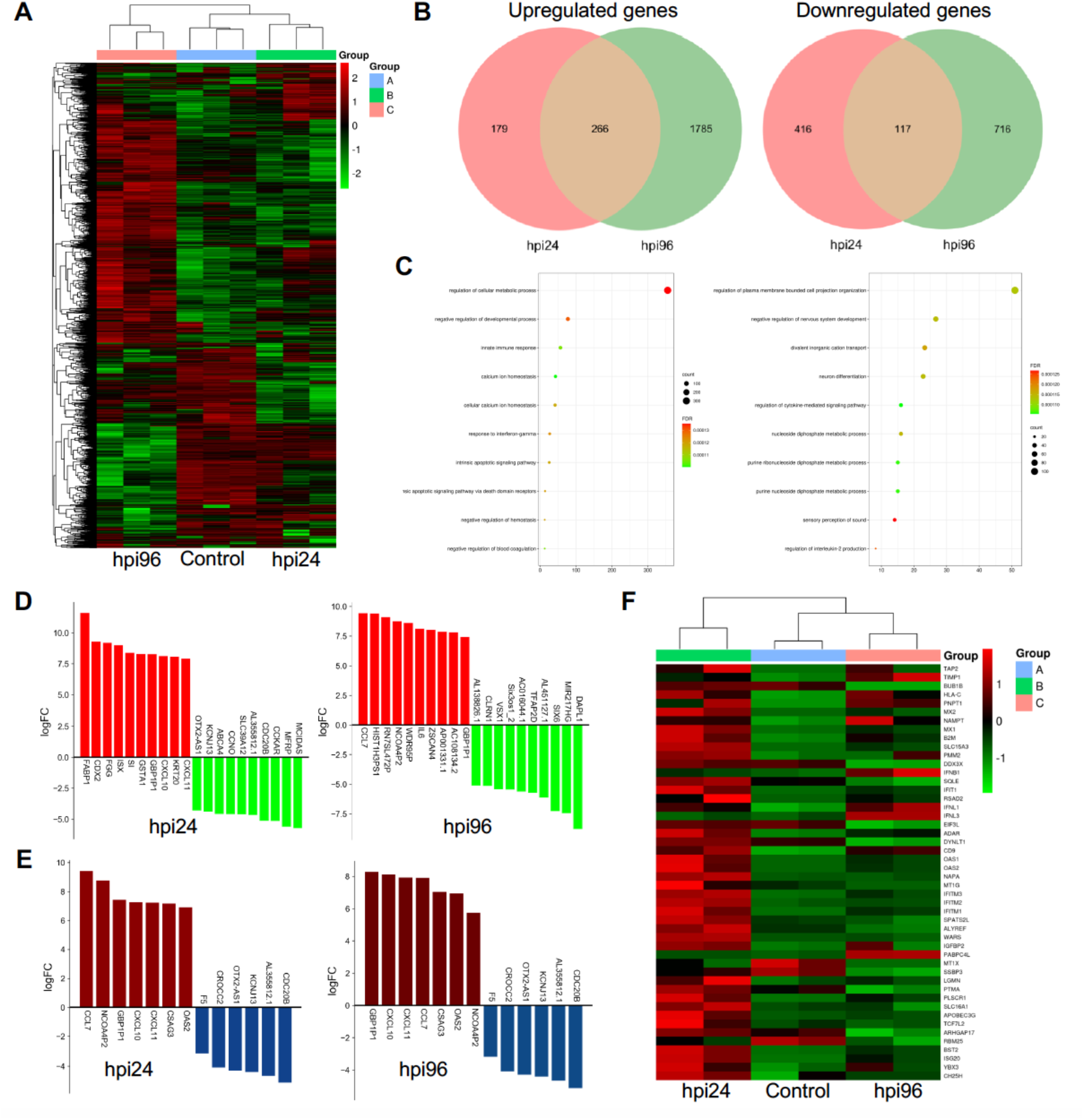
RNA transcriptomic analysis of human brain organoids after WSN infection. (**A**) Hierarchical clustering heatmap of differentially expressed genes derived from a comparison of a group of control, hpi 96, and hpi 24. (**B**) Venn diagram of upregulated and downregulated genes of brain organoids infected with WSN at 24 hpi and 96 hpi. (**C**) Top 10 enriched GO terms in brain organoids infected with WSN at 24 hpi and 96 hpi. (**D**) Top 10 upregulated and downregulated genes at 24 hpi and 96 hpi after WSN infection. (**E**) The upregulated and downregulated genes at 24 hpi and 96 hpi after WSN infection. (**F**) Heatmap of interferon stimulating genes (ISGs).

### WNS impairs brain organoid growth through inducing apoptosis and inflammation

To further investigate the neuropathogenesis of brain organoids subjected to WSN infection, we conducted related assays, including apoptosis and inflammatory factor release after WSN infection at 1 dpi and 4 dpi (**Fig. 5A**). Significant cell apoptosis of SOX2+ NSCs and MAP2+ neurons in day 30 and day 60 brain organoids at 1 dpi and 4 dpi was observed by TUNEL staining (**Fig. 5B, C**). The percentage of TUNEL+ cells was a time-dependent increase, and the apoptosis of NSCs was higher than neurons during early WSN infection (**Fig. 5D, E**). Unlike Zika virus infection, which mainly induces the apoptosis of NSCs to result in microcephaly(Tang et al., 2016), WSN primarily infects MAP2+ neurons and causes the apoptosis of both NSCs and neurons. We also detected the apoptosis of GFAP+ astrocytes in day 250 organoids, and we did not observe obvious apoptosis (**Fig. 5F**). Next, we determined the inflammatory factor levels in supernatants from days 30 and 60 brain organoids by enzyme linked immunosorbent assay (ELISA). We found that TNF-α, INF-γ, IL-6, CCL2, and COX2 in organoid supernatants were significantly increased in a time-dependent manner from days 30 and 60 brain organoids (**Fig. 5G, H**). At the cellular level, we also confirmed that influenza virus could induce NSC apoptosis (**Fig. S1C, D)** and inhibit proliferation (**Fig. S1E, F)**, but did not induce astrocyte apoptosis (**Fig. S1I)**. Taken together, these results suggested that WSN could impair brain organoid growth through eliciting apoptosis and inflammation (**Fig. 5I**).

**Figure 5.**
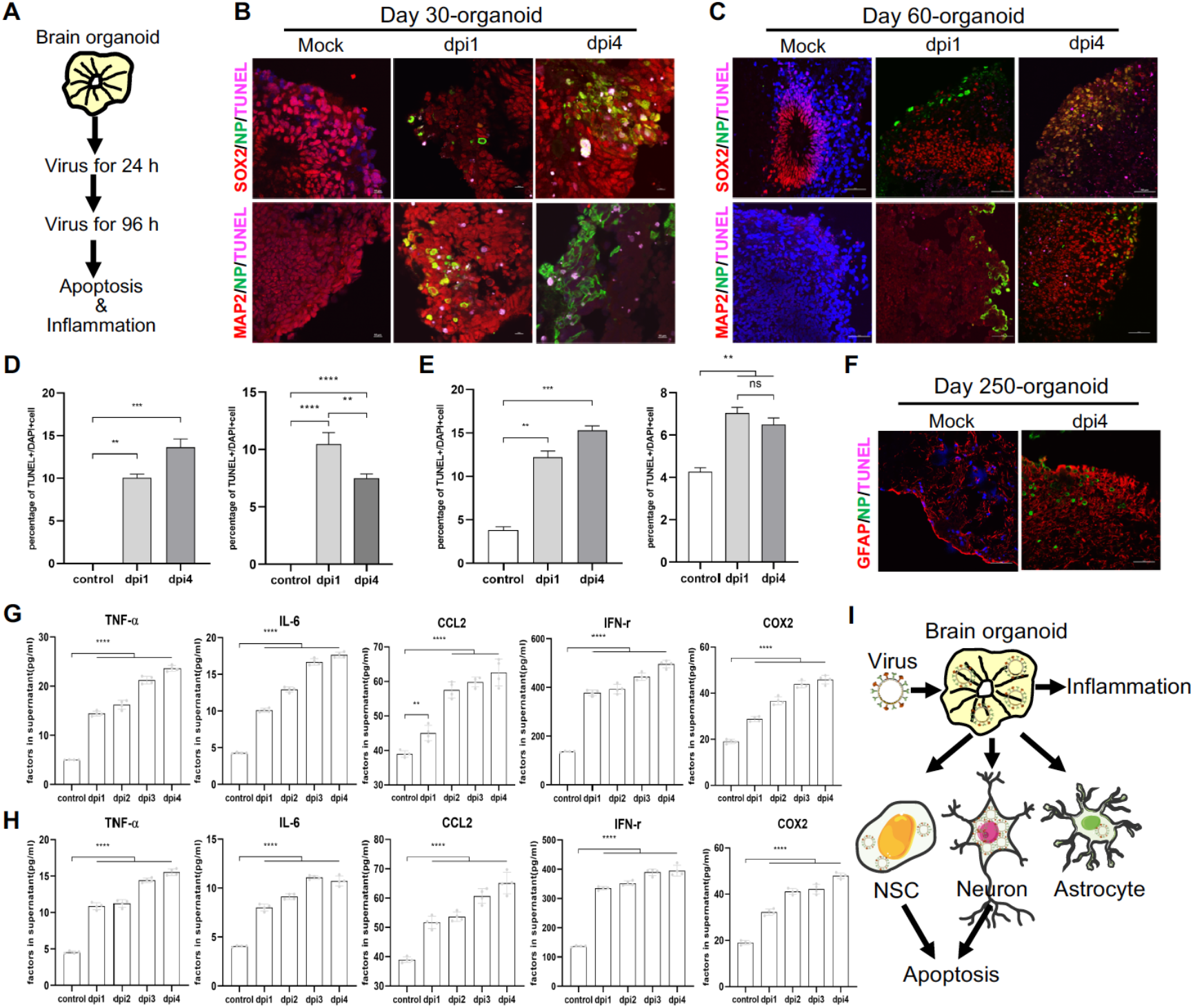
WSN induced cell apoptosis and inflammation in human brain organoids. (**A–D**) TUNEL staining and quantification of positive cells in day-30 and day-60 brain organoids infected with WSN at 1 dpi and 4 dpi show a time-dependent increase, respectively. Scale bars, 10 µm. (**E**) TUNEL staining of GFAP+ astrocytes in day 250 brain organoids at 4 dpi. (**F, G**) Secreted inflammatory factors (e.g., TNF-α, IL-6, CCL2, IFN-γ, and COX2) in day-30 (**F**) and day-60 (**G**) brain organoids at indicated infection timepoints. Scale bars, 100 µm.

### Antiviral and neuroprotective effects of drugs and neurotrophic factors

As these results demonstrated, WSN did impair the human brain through inducing cell apoptosis and inflammation responses using *in vitro* brain organoid models. Therefore, we performed antiviral screening studies. We first analyzed several compounds, PYC-12, RO3306, and WHL-50B, at the cellular level, and we used nucleozin as a positive control, as it induces nuclear accumulation of influenza virus nucleoprotein (NP) leading to cessation of viral replication (Gerritz et al., 2011; Kao et al., 2010). Our results showed that PYC-12, WHL-50B, and nucleozin all significantly suppressed virus infection compared to RO3306 (**Fig. S4A**), and PYC-12 could also suppress WSN infection in astrocytes, but not as clearly as nucleozin (**Fig. S4B**). Next, we further evaluated the antiviral effect of four compounds at the organoid level. Briefly, day 40 brain organoids were first treated with four compounds for 2 hours, followed by co-treatment with WSN and compounds for 1 hour, followed by continued compound treatment for the indicated number of days (**Fig. 6A**). Representative bright field images showed that PYC-12 could significantly rescue the morphological changes of brain organoids compared to the WSN infection group (**Fig. 6B**). Additionally, three compounds, peramivir, PYC-12, and WHL-50B, were used in drug screening of H3N2-infected organoids, and the results showed that peramivir as a positive drug, which is a highly selective inhibitor of influenza A and B neuraminidase (Kohno et al., 2011), had the highest antiviral effects compared to other drugs (**Fig. S5**). Statistical analysis of the diameter and area of organoids treated with PYC-12 also demonstrated its rescue effects (**Fig. 6C, D**). Therefore, we used PYC-12 for our following studies. Immunostaining of SOX2+ NSCs and MAP2+ neurons on day 30 and day 60 brain organoids infected by WSN revealed that PYC-12 treatment could significantly inhibit WSN infection (**Fig. 6E**) and cell apoptosis (**Fig. 6F**) at 1 dpi and 4 dpi. PYC-12 also decreased the extracellular and intracellular viral titers (**Fig. 6G**), as well as decreasing IL-6 and TNF-α production (**Fig. 6H**). Microelectrode array (MEA) analysis showed that PYC-12 could increase the weighted mean firing rate (Hz) of brain organoids compared to WSN infection at indicated time points (**Fig. 6I, J**). Collectively, we screened a potential drug candidate-PYC-12 against WSN infection using brain organoid models (**Fig. 6K**).

**Figure 6.**
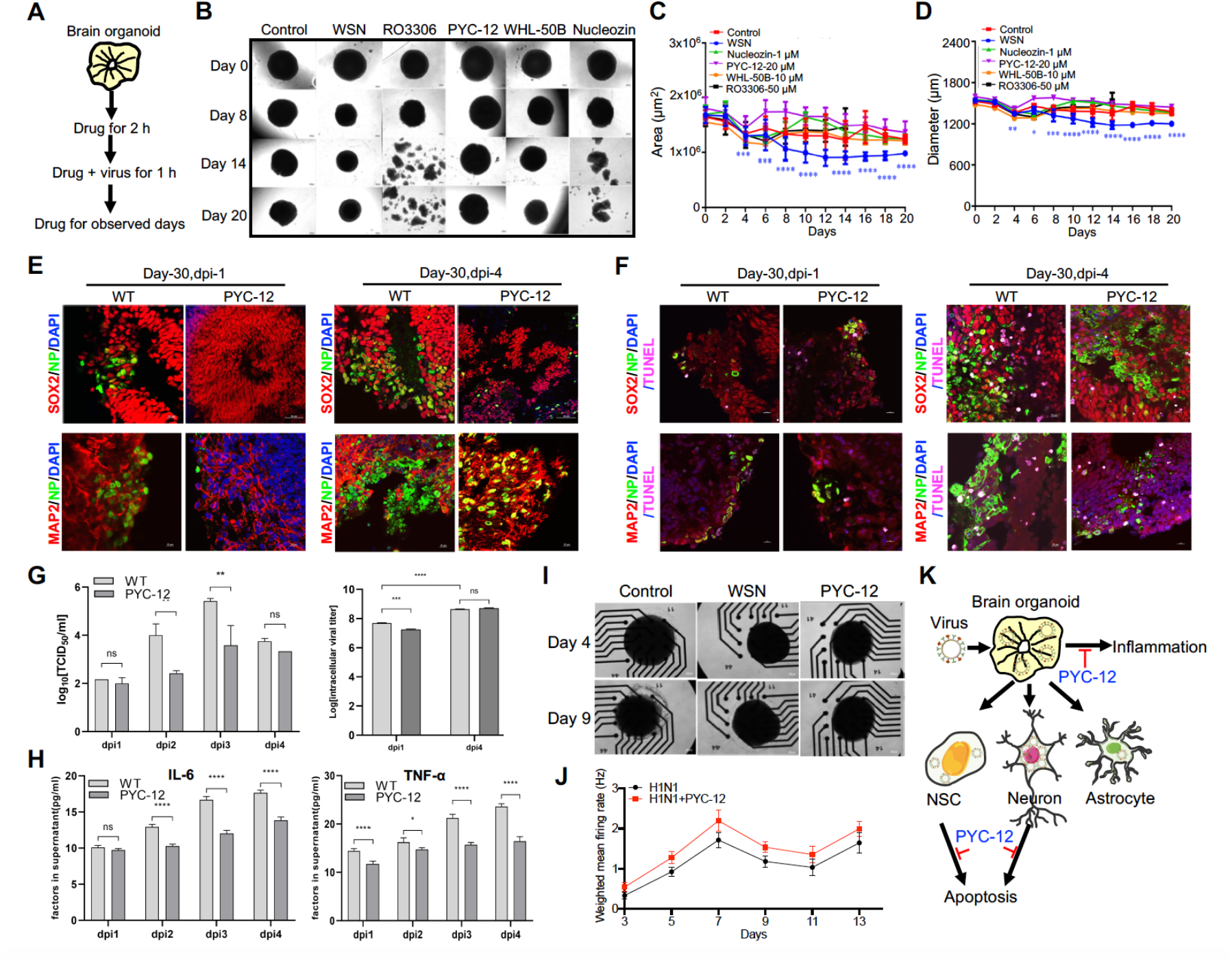
Antiviral drug study of human brain organoids infected with influenza virus. (**A**) Schematic of the workflow for drug screening. (**B**) Representative bright field images of organoids cotreated with H1N1-WSN and several drugs at indicated time points. Nucleozin was used as a positive control. Scale bars, 50 µm. (**C, D**) Statistical analysis of area (µm^2^) and diameter (µm) of brain organoids infected with H1N1-WSN at indicated time points. (**E**) Immunostaining of neural stem cells and neurons in day 30 brain organoids co-treated with WSN and PYC-12 for 1 day and 4 days. Scale bars, 10 µm and 50 µm. (**F**) TUNEL staining of neural stem cells and neurons in day 30 brain organoids co-treated with WSN and PYC-12 for 1 day and 4 days. Scale bars, 10 µm. (**G**) The viral titers of intracellular and supernatants from day 30 brain organoids co-treated with WSN and PYC-12. (**H**) The inflammatory factors release (IL-6 and TNF-α) of day-30 brain organoids co-treated with WSN and PYC-12 at indicated infection timepoints. (**I**) The representative bright field images of microelectrode array (MEA) analysis of PYC-12 treated brain organoids. Scale bars, 50 µm. (**J**) Weighted mean firing rates (Hz) of brain organoids treated with H1N1 or cotreated with H1N1 and PYC-12 at indicated time points. (**K**) Schematic illustration of the antiviral strategy of PYC-12 through anti-apoptosis and anti-inflammation.

In addition, neurotrophic factors (NFs), which are endogenous soluble proteins regulating the survival and growth of neurons, protect human normal brain functions against microbial pathogens have been reported in the literature (Platholi and Lee, 2018). Thus, to investigate whether NFs also have antiviral effects in *in vitro* organoid models, we conducted antiviral assays with the same treatments (**Fig. 6A**). First, we demonstrated that brain-derived neurotrophic factor (BDNF), glial-derived neurotrophic factor (GDNF), neurotrophin-3 (NT3), and their combinations (GBN) could significantly inhibit WSN infection at the NESTIN+ NSCs level (**Fig. S6A**), especially BDNF and GBN in NP+ cells (**Fig. S6B**). All three NFs and GBN showed decreased intracellular virus titers (**Fig. S6C**), while only GBN inhibited extracellular virus titers at the NSC level (**Fig. S6D**). Then, we confirmed the antiviral effect of GBN on day 30 organoids (**Fig. S6E, F**), and GBN showed decreases in intracellular and extracellular virus titers (**Fig. S6G, H**). GBN significantly inhibited WSN infection through decreasing both cell apoptosis (**Fig. S6I, J**) and inflammatory factor release (**Fig. S6K**). At the gene level, we found that NF could significantly induce a set of ISG expressions, such as *MCL1, IFITM3, B2M, BST2, OAS1*, and *PKR* to function against virus infection (**Fig. S7**). Collectively, neurotrophic factors also possessed antiviral roles via inducing ISG expression to suppress cell apoptosis and inflammation response in brain organoid models (**Fig. S6L**), and implicating the possibility of a combination therapy in the future.

## Discussion

The neuropathogenesis of the human brain caused by influenza virus remains poorly understood. In the light of this fundamental problem, we designed an experimental platform using human brain organoids as a virus infection model to study this neuropathogenesis in detail. Brain organoids serve as an invaluable tool in the field of virus research to investigate the molecular events underlying virus replication and antiviral drug screening.

In this study, we first demonstrated that influenza virus (H1N1-WSN) could widely infect multiple cell types in brain organoids, including SOX2+ neural stem cells (NSCs), MAP2+ neurons, and GFAP+ astrocytes, and holds an infection tropism of MAP2+ neurons (**Fig. 3**), which may support a direct link between influenza virus infection and the neurologic symptoms. Inflammatory factors (e.g., IL-6, TNF-α, and INF-γ) induced by WSN infection in brain organoids (**Fig. 5G, H**) may be the major cause for viral entrance into the brain through damaging the blood brain barrier (BBB) (Shlosberg et al., 2010). However, whether other virulent influenza viruses (e.g., H3N2, H5N1, or H7N9) have similar results needs further investigation in higher level of biosafety laboratories. Unlike WSN infection, Zika virus preferentially infects human cortical neural progenitor cells with high efficiency, but exhibits lower levels of infection in human embryonic stem cells (ESCs), hPSCs, and immature cortical neurons (Wu et al., 2018). Therefore, regional and cell type specific tropism of virus infection may directly support a link with specific pathological syndromes.

As is well known, although brain organoids can serve as a model of a three-dimensional organ to achieve multiple cellular interactions not possible with cell lines, organoids are very immature and only partly model the human brain. Therefore, which type of neurons are specifically infected with influenza virus in brain organoids needs further study in our next work. Some research groups have leveraged virulent viruses (e.g., H5N1, H5N3, or H3N2) (Mori and Kimura, 2001) to infect mice, and have demonstrated the route of brain infection, but cell type-specific tropism of viral infection in the central nervous system remain poorly understood. Therefore, if we completely confirm the cell type-specific infections of influenza or other viruses, it should provide a scientific basis for developing specific neurological disorder drugs. Single-cell RNA sequencing technology (Hwang et al., 2018; Ofengeim et al., 2017) could be used to track specific cell types infected by viruses in space and time through detecting the gene expression of viruses, while human brain tissue infected by viruses are very hard to get and labeling hundreds or thousands of nerve cells is difficult for current single-cell RNA sequencing technology.

In addition, we found that WSN could infect astrocytes (**Fig. 3J**), but not induce their apoptosis (**Fig. 5F, S1**), which was similar to brain organoids infected with SARS-CoV-2 (Song et al., 2020). This finding may indicate that virus-infected astrocytes can lead to peripheral cell death by creating a locally hypoxic and a resource-restricted environment for cells. Furthermore, we performed whole RNA transcriptomic analysis for comparison of a control and 4 dpi and 1 dpi group and identified some upregulated protein targets (e.g., CCL7, NCOA4P2, GBP1P1, CXCL10, CXCL11, CSAG3, and OAS2) and noncoding RNAs (e.g., BISPR, AC116407.2, AC092687.3, and MIR4435-2HG), implicating these for the development of new antiviral targets. However, whether these targets can exert antiviral effects needs to be further validated in future work. Finally, we performed antiviral screening using brain organoids and found a potential antiviral drug, PYC-12, which could significantly suppress virus replication, apoptosis of NSCs and neurons, and inflammatory responses (**Fig. 6**). However, whether the identified drug could enter into the brain through the BBB needs to be validated in animal models. Next, we intend to conduct a high throughput antiviral drug screening study using this organoid platform. Moreover, we also explored the antiviral and neuroprotective effects of neurotrophic factors such as GDNF, BDNF, and NF3 (**Fig. S6**), and demonstrated that they also did suppress WSN infection, the apoptosis of NSCs and neurons, and inflammatory responses. This work also highlights the possibility of using a combinations therapy of drugs and neurotrophic factors in clinical treatment for viral infection.

## Conclusion

In this study, using an *in vitro* brain organoid model, we demonstrated that MAP2+ neurons were more susceptible to influenza virus (H1N1-WSN) infection, unveiled the neuropathogenesis of the brain through inducing apoptosis and inflammation, and conducted antiviral screening to achieve antiviral and neuroprotective aims. In summary, we establish a tractable experimental model system to investigate the impact and mechanism of influenza virus on human brain development, and provide a platform for identifying therapeutic compounds.

## Materials and Methods

### Cell culture and reagents

Human HEK293T (CRL-11268) and Madin-Darby Canine Kidney (MDCK) cells (ATCC, CRL-2936) were maintained in Dulbecco’s Minimal Essential Medium (DMEM) (Gibco) supplemented with 10% fetal bovine serum (Gibco), 100 units/mL penicillin, and 100 µg/mL streptomycin (Invitrogen). Authentication and test for the free of mycoplasma were performed with MycAway™ one-step mycoplasma detection kit (Yeasen). Astrocytes were purchased from Cellapy (CA2315106) and cultured in NeuroEasy maintenance medium (Cellapy). Human embryonic stem cells (hESCs) were obtained from Harvard Stem Cell Institute. hESCs were routinely checked for pluripotent, normal karyotype, mycoplasma free and cultured in feeder-free conditions on Matrigel-coated plates with Essential 8 medium (GIBCO) and passaged with TrypLE™ express (GIBCO). 10 µM Peramivir (M3222, AbMole BioScience), 1 µM Nucleozin (A3670, Apexbio), 50 µM PYC-12, 20 µM RO3306, 10 µM WHL-50B, 20 ng/mL BDNF (450-02, PeproTech), 20 ng/mL GDNF (450-10, PeproTech), 20 ng/mL NT3 (450-03, PeproTech) were used in this study.

### Viruses

Three H1N1 strains (WSN, CA07 and PR8), Enterovirus 68/71, and SFTSV obtained from Academy of Military Medical Sciences were used in this study. All viruses’ stocks were prepared in MDCK cells and titrated by TCID_50_ on MDCK cells as described in details below. Studies with infectious H1N1 were conducted under biosafety level 2 (BSL-2) conditions at the Peking University Health Science Center with approval from Institutional Biosafety Committee.

### Neural stem cells (NSCs) differentiation

hESCs were differentiated into NSCs on Matrigel-coated plates using the monolayer protocol as previously described(Lippmann et al., 2014). Briefly, hESCs were first dissociated into single cells with Accutase (STEMCELL). Then, hESCs were plated onto Matrigel at a density of 2×10^5^ cells/cm^2^ in E8 medium containing 10 µM ROCK inhibitor (STEMCELL Technologies) and cultured overnight. 24 h later, cells were changed to E6 medium (GIBCO) containing 10 µM SB431542 (STEMCELL) and 100 nM LDN193189 (Selleckchem) to initiate differentiation. Medium was changed every day until day 7. Day 7-NSCs were used for the following experiments.

### Brain organoids generation and culture

Brain organoids were generated from hESCs as previously described(Lancaster and Knoblich, 2014), but slightly modified. Briefly, for embryoid body (EB) formation, hESCs were washed twice with DPBS, incubated with Accutase for 5 minutes, and dissociated into single cells. 3000 single cells were seeded in each well of low attachment 96-well U-bottom plate in E8 medium containing 10 µM ROCK inhibitor and centrifuged at 100 g for 3 min, then medium was half changed every other day. On day 4, EBs were transferred to low attachment 24-well plate in neural induction medium containing DMEM-F12 (GIBCO) with 1% N2 supplement (GIBCO), 1% Glutamax supplement (GIBCO), 1% MEM-NEAA (GIBCO) and 1 µg/mL Heparin (GIBCO), and medium was changed after 48 h. On day 7, EBs were transferred into Matrigel droplets as previously described and cultured in brain organoid differentiation media containing 50% DMEM-F12, 50% Neurobasal, 200x N2 supplement, 0.025% Insulin (GIBCO), 100x Glutamax supplement, 200x MEM-NEAA, 100x penicillin-streptomycin, 0.035% 2-Mercaptoethanol and 100x B27 supplement without Vitamin A, and medium was changed after 48 h. On day 10, organoids were transferred to orbital shaker (Corning) in brain organoid differentiation media with Vitamin A, medium was changed every 4 days.

### Virus infection

For cell line infection, H9, NSCs and astrocytes were seeded in chamber at 1×10^6^ cells. The cells were then rinsed with PBS, and WSN was diluted to the desired multiplicity of infection (MOI) of 1 and added to the cells. The cells were incubated for 2 h at 37 °C. The supernatant was removed and the cells were washed twice with PBS. Culture medium with 1% FBS and 1000x TPCK (final concentration 1 µg/mL) was added to each well, and cells were incubated at 37 °C and 5% CO_2_, after 24 h infection, cells were prepared for immunofluorescence staining. For brain organoids infection, organoids were transferred to low attachment 24-well plate and washed twice with DPBS, and WSN was diluted to the desired multiplicity of infection (MOI) of 1 and added to the cultured medium. The organoids were incubated for 8 h at 37 °C. The supernatant was removed and the cells were washed twice with DPBS, and then organoids were transferred to low attachment 6-well plate, 4 mL culture medium with 1% FBS and 1000x TPCK (final concentration 1 µg/mL) was added to each well, and organoids were incubated at 37 °C and 5% CO_2_ at a shake speed of 60 rpm, after 24 h and 96 h infection, organoids were prepared for immunofluorescence staining, RNA extraction and RNA-seq, and the supernatant was harvested for ELISA and virus titration.

### Immunofluorescence staining

For cell immunofluorescence staining, the cells were fixed with 4% PFA at room temperature for 10 min, permeated with PBST (PBS with 0.1% Triton X-100) for 20 min and blocked with 1% BSA for 30 min. Then the cells were incubated with primary antibodies listed in **Supplementary Table 1** at 4 °C overnight. The cells were subsequently incubated with secondary antibodies listed in **Supplementary Table 1** at room temperature for 1 h. The cells were mounted with mounting fluid containing DAPI (Yeason, 36308ES11). For organoid immunofluorescence staining, the organoids were fixed with 4% PFA at room temperature for 30 min, then immersed in 30% (w/v) sucrose until submersion before embedding and freezing in the Optimal Cutting Temperature (OCT) compound (Tissue-Tek). Serial 12 µm sections were obtained by cryo-sectioning of the embedded organoid at −20 °C using a cryostat (Leica). Cryosections were permeated with in PBST at temperature for 30 min and blocked with sheep serum (Zhongshanjinqiao, ZLI-9022) for 1 h. The sections were incubated with primary antibodies listed in **Supplementary Table 1** diluted in blocking buffer at 4 °C overnight. The slides were subsequently incubated with secondary antibody at room temperature for 1 h. The slides were mounted with mounting fluid containing DAPI. Stained sections were photographed under a Nikon Ti-S microscope. Apoptotic cells were labelled with Click-It Plus TUNEL assay (C10619, ThermoFisher Scientific).

### RNA isolation and quantitative RT-PCR

Total RNA was extracted from cells or organoids using the Quick-RNA MicroPrep kit (Zymo Research). RNA was subjected to quantitative real-time PCR in accordance with the protocol provided by one-step SYBR green RT-PCR Kit (Cwbio). The transcripts were quantitated and normalized to the internal GAPDH control. The primers used in the experiments are listed in **Supplementary Table 2**. The PCR conditions were 1 cycle at 95 °C for 5 min, followed by 40 cycles at 95 °C for 15 s, 60 °C for 1 min, and 1 cycle at 95 °C for 15 s, 60 °C for 15 s, 95 °C for 15 s. The results were calculated using the 2^-ΔΔCT^ method according to the GoTaq qPCR Master Mix (Promega) manufacturer’s specifications.

### Virus quantification by 50% tissue culture infective dose

For quantifying all viruses’ stocks, the 50% tissue culture infectious dose (TCID_50_/mL) titers were determined. In brief, 5×10^4^ HEK293T cells were seeded in 96-well plates the day before infection. The virus samples were serially diluted with DMEM containing 1% FBS (10^3^ to 10^10^) and then each of dilution was added in wells separately. The plates were incubated at 37 °C in 5% CO_2_ for 2-5 days. The cytopathic effect (CPE) was observed under a microscope and determined virus titer using the Reed-Münch endpoint calculation method.

### Microelectrode arrays (MEA)

Day 40 brain organoids were seeded onto 48-well transparent MEA plates. Brain organoids were cultured in brain organoid differentiation media containing 50% DMEM-F12, 50% Neurobasal, 200x N2 supplement, 0.025% Insulin (GIBCO), 100x Glutamax supplement, 200x MEM-NEAA, 100x penicillin-streptomycin, 0.035% 2-Mercaptoethanol and 100x B27 supplement with Retinoic Acid (RA). MEA recordings were performed on day 3, 5, 7, 9, 11, 13 at 37 °C in a Maestro MEA system with AxIS software using a bandwidth with a filter for 10Hz to 2.5 kHz cutoff frequencies. For the pharmacological experiment, 50 µM PYC-12 were applied to plate immediately before recording. For MEA recording, brain organoids treated with WSN were included as the control organoids. The phase contrast images of organoids seeded in the MEA plates were taken after MEA recording.

### Enzyme-linked immunosorbent assay (ELISA)

Inflammatory factors (TNF-α, INF-γ, IL-6, CCL2, COX2) in cultured supernatants of brain organoids with or without challenging by virus was measured using a commercial ELISA Kit (Dogesce). Briefly, samples were double diluted using the dilution buffer, and the optical density (OD) was measured at 450 nm with an ELISA reader (Beckman). The concentration of inflammatory factors was calculated according to the manufacturer’s instruction.

### The whole transcriptome analysis

High throughput RNA sequencing was performed by Cloud-Seq Biotech (Shanghai, China). Total RNA was extracted from organoids (three biological replicates for each group) by TRIzol and the rRNAs were removed with NEBNext rRNA Depletion Kit (New England Biolabs, Inc., Massachusetts, USA). RNA libraries were constructed using the NEBNext® Ultra(tm) II Directional RNA Library Prep Kit (New England Biolabs, Inc., Massachusetts, USA) following the manufacturer’s instructions. The libraries were quality-controlled and quantified using the BioAnalyzer 2100 system (Agilent Technologies, Inc., USA). The library sequencing was performed on an Illumina Hiseq instrument with 150 bp paired end reads. Paired-end reads were harvested from Illumina HiSeq 4000 sequencer, and quality-controlled by Q30. After 3’ adaptor-trimming and removing low-quality reads by cutadapt software (v1.9.3), high-quality clean reads were aligned to the reference genome (UCSC MM10) with hisat2 software (v2.0.4). Guided by the Ensembl gtf gene annotation file, the cuffdiff software (part of cufflinks) was used to obtain the gene level FPKM as the expression profiles of mRNA. The total expressed gene number and LogFPKM of mRNA in different mouse groups were plotted and compared.

### Statistical analysis

All data were analyzed using the GraphPad Prism 9 software. For the statistical analysis of other results, statistical evaluation was performed by Student’s unpaired t-test or one-way ANOVA with Tukey’s multiple comparisons test. Data are presented as means ± SD or as described in the corresponding legends. A probability of p < 0.05 was considered as statistically significant. For annotations of significance, ^*^p<0.05; ^**^p<0.01; ^***^p<0.001; ^****^p<0.0001.

## Data availability

All other data supporting this study are available within this paper and its Supplementary Information.

## Acknowledgements

This work was funded by China’s National Science and Technology Major Projects for Major New Drugs Innovation and Development (No. 2018ZX09711003-001-003).

## Author contributions

Q. X. conceived the idea and X. Z. initiated the project. X. Z. and H. L. analyzed the sequencing data, prepared the figures, and wrote the draft. Z. X. performed the flow cytometry and cytokine release activity assays. All authors had access to the data. K. Y., L. T. and L. D. verified the data. All co-first authors and corresponding authors discussed the results and revised the paper.

## Additional information

**Supporting information** is available for this paper.

## Supporting information

**Figure S1.**
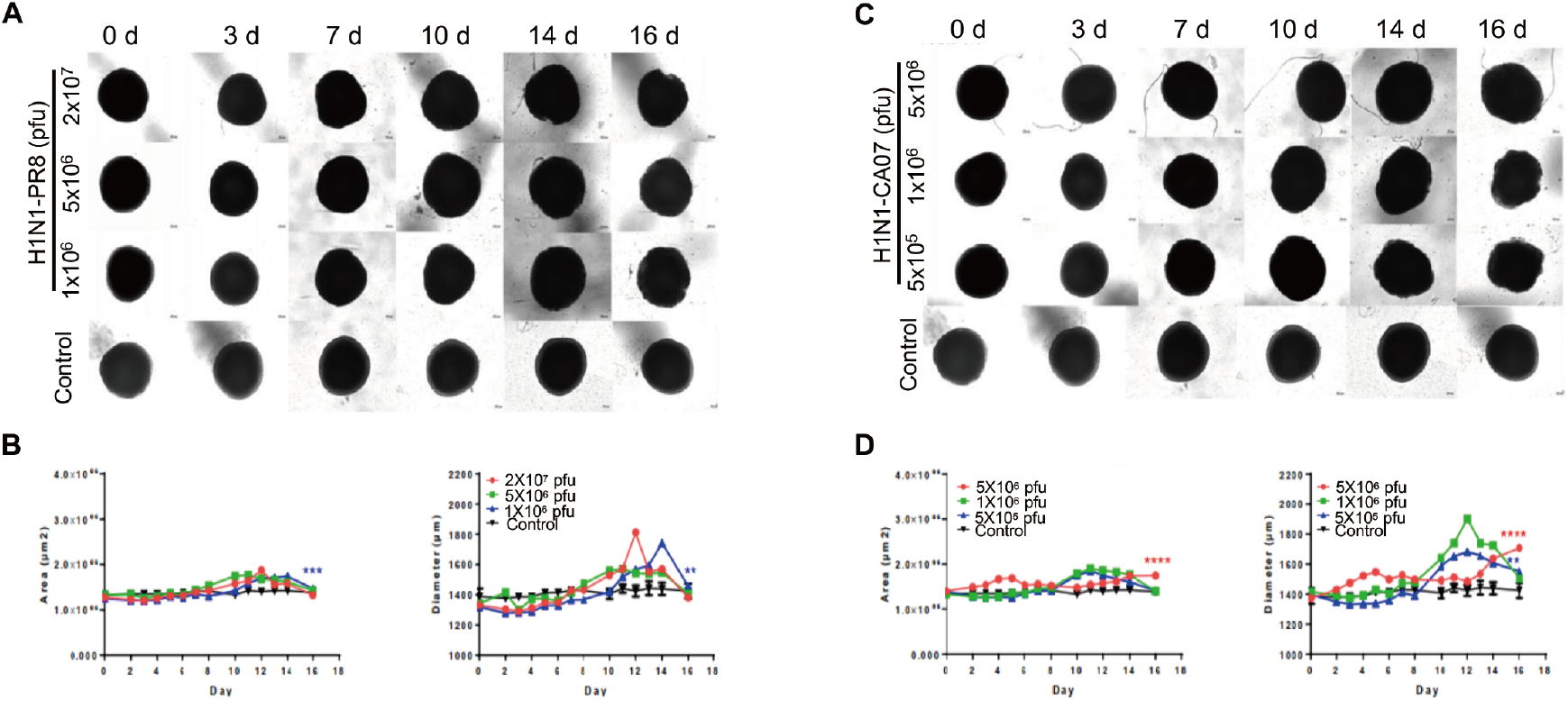
(**A, C**) Representative bright-field images of day 40-brain organoids infected with H1N1-PR8 or H1N1-CA07 at indicated time points. The infected concentrations for H1N1-PR8 were 1×10^6^ pfu, 5×10^6^ pfu, and 2×10^7^ pfu; for H1N1-CA07, they were 5×10^5^ pfu, 1×10^6^ pfu, and 5×10^6^ pfu. Scale bars, 50 µm. (**B, D**) Statistical analysis of area (µm^2^) and diameter (µm) of brain organoids infected with viruses at indicated time points.

**Figure S2.**
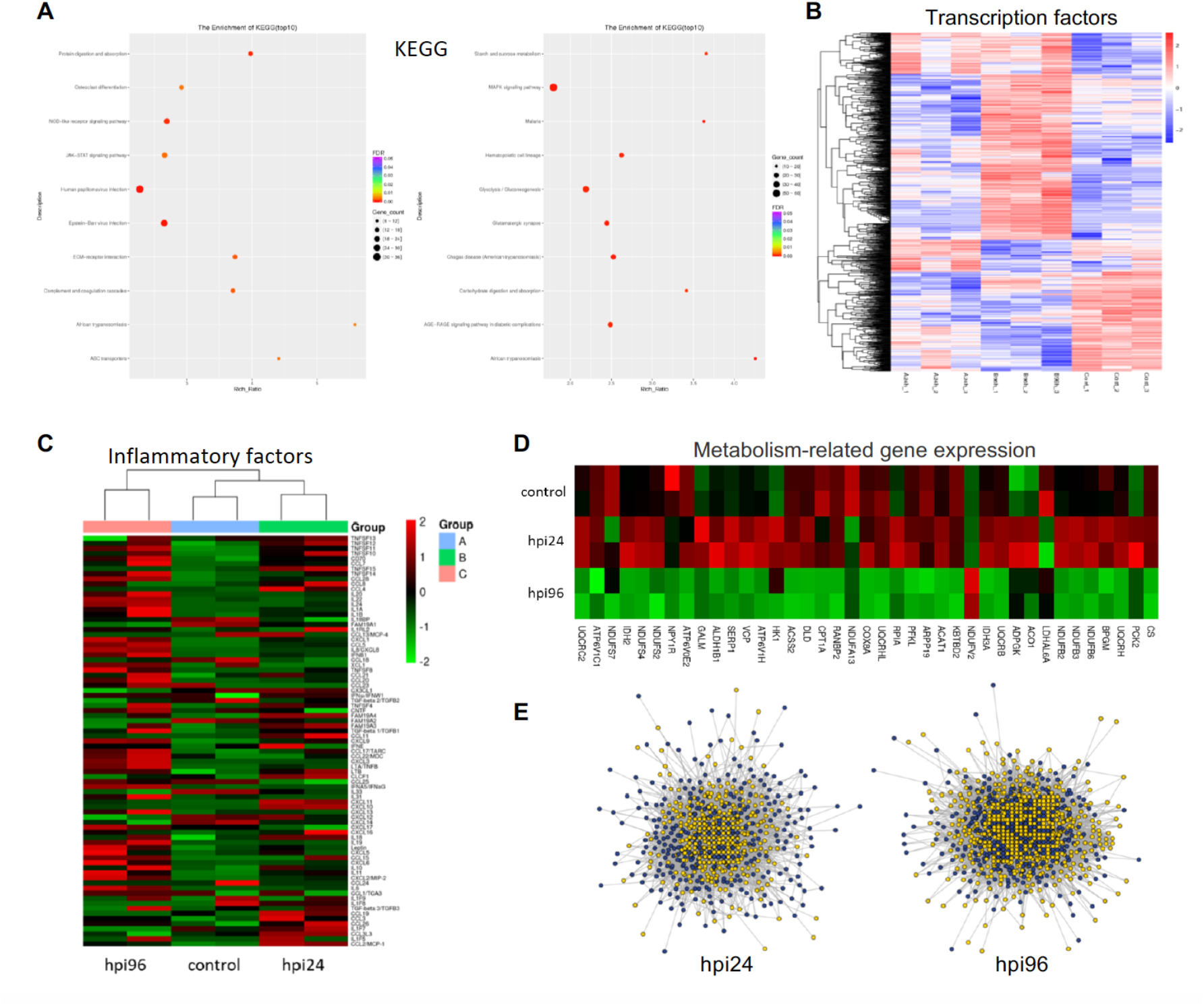
(**A**) The top 10 KEGG pathway enrichments of of brain organoids infected with WSN at 1 dpi and 4 dpi. (**B**) Heatmap of transcriptional factor expression in control, hpi24, and hpi96 groups. (**C**) Heatmap of inflammatory factor expression in control, hpi24, and hpi96 groups. (**D**) Heatmap of metabolic genes. (**E**) Protein-protein interaction network of WSN infected brain organoids at 24 hpi and 94 hpi.

**Figure S3.**
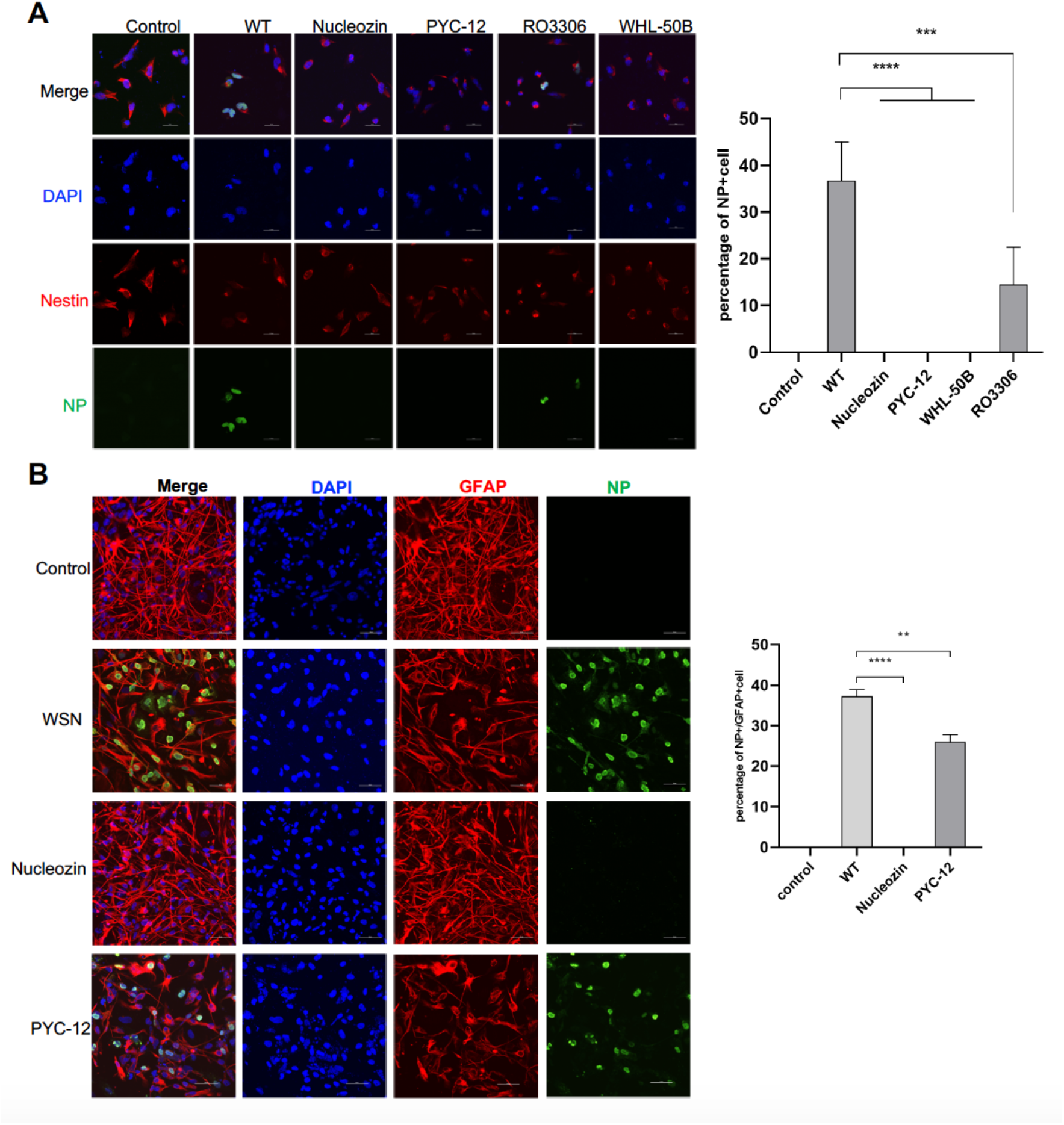
Antiviral drug screening using NSCs and astrocytes. (**A**) Immunostaining and statistical analysis of neural stem cells treated with WSN, Nucleozin, PYC-12, RO3306, and WHL-50B. Non-treated NSCs were used as a negative control. Scale bars, 20 µm. (**B**) Immunostaining and statistical analysis of astrocytes treated with WSN, nucleozin, and PYC-12. Non-treated astrocytes were used as a negative control. Scale bars, 50 µm.

**Figure S4.**
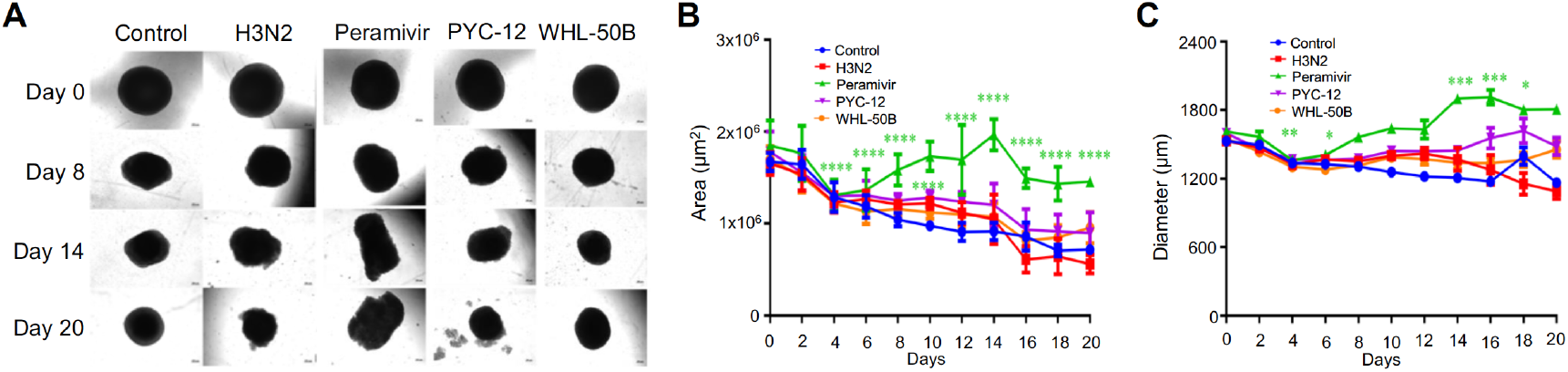
Antiviral drug study of human brain organoids infected with H3N2-HKT68. (**A**) Phase-contrast images of brain organoids cotreated with H3N2-HKT68, peramivir, PYC-12, and WHL-50B at indicated time points. Scale bars, 50 µm. (**B, C**) Statistical analysis of the area (µm^2^) and diameter (µm) of organoids cotreated with H3N2-HKT68 and different drugs at indicated time points.

**Figure S5.**
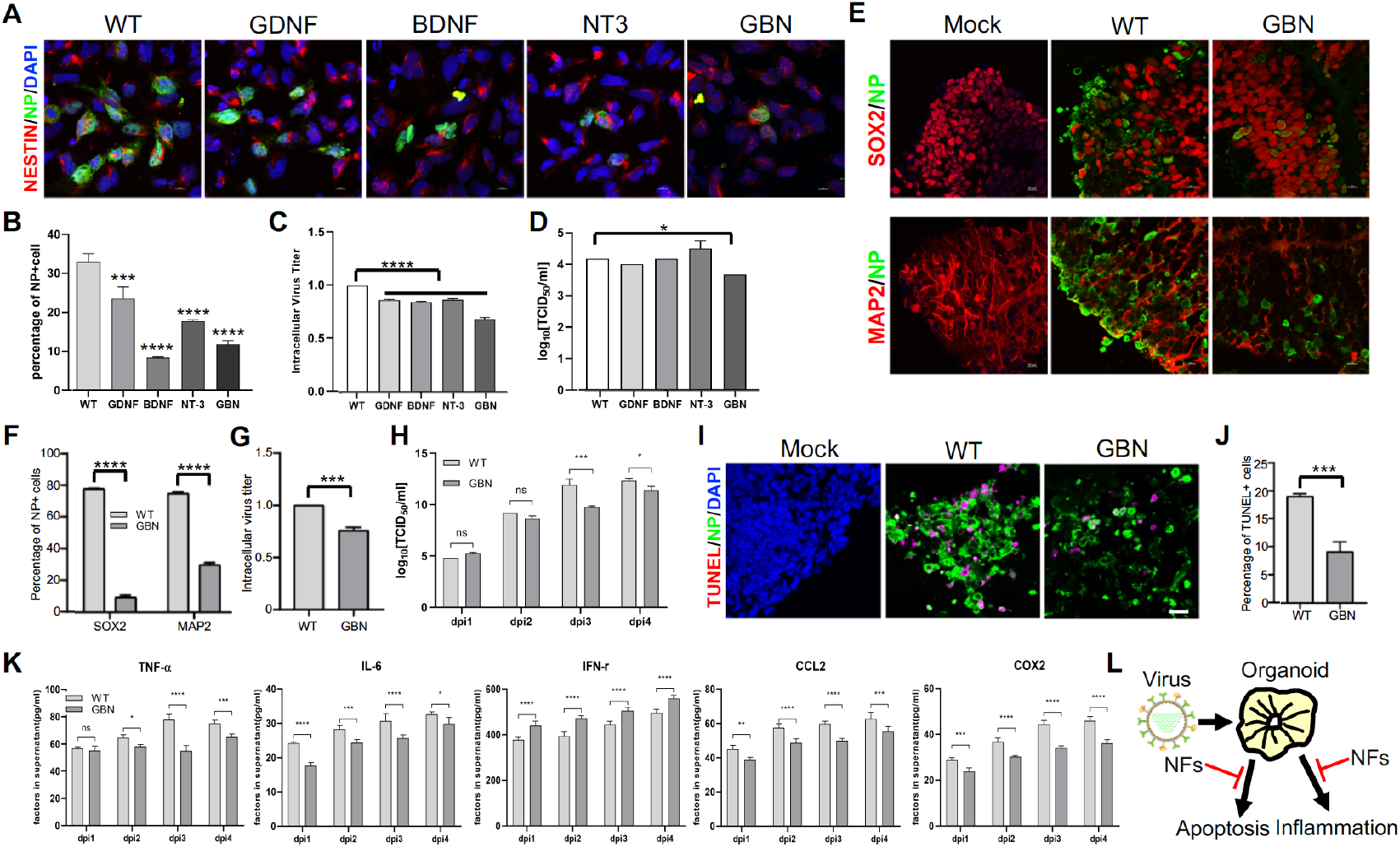
Neurotrophic factors inhibits WSN infection. (**A**) Immunostaining of NSCs treated with WSN or different neurotrophic factors, including BDNF, GDNF and NT3. GBN denotes the combination of these three neurotrophic factors. Scale bar, 10 µm. (**B**) The percentage of NP+ cells after treatment. (**C, D**) Intracellular (left) and extracellular (right) virus titers after treatment. (**E**) Immunostaining of day 30 brain organoids treated with WSN and GBN. Scale bar, 10 µm. (**F**) The percentage of NP+ cells after GBN treatment compared to WSN infection. (**G, H**) The intracellular and extracellular virus titers after treatment. (**I, J**) TUNEL staining and quantification of positive cells in day 30 brain organoids at 4 dpi. Scale bar, 50 µm. (**K**) The secreted inflammatory factors (e.g., TNF-α, IL-6, CCL2, IFN-γ, and COX2) from day 30 brain organoids at indicated infection timepoints. (**L**) Schematic illustration of antiviral methods produced through neurotrophic factors treatment.

**Figure S6.**
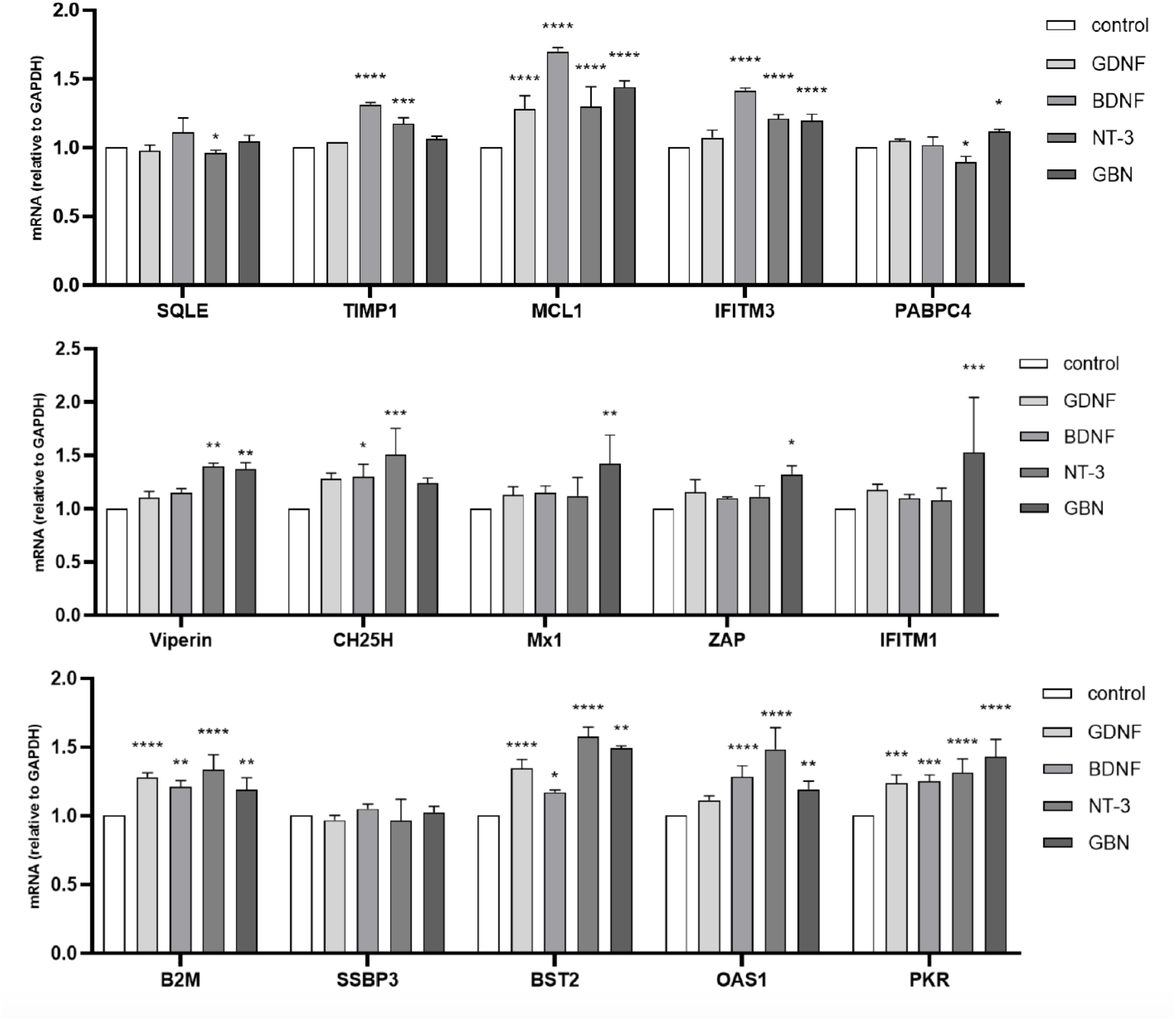
Insulin stimulating gene expression from organoids cotreated with WSN and neurotrophic factors by quantitative RT-PCR.

